# Free radical production induced by visible light in live fruit flies

**DOI:** 10.1101/2024.02.19.580981

**Authors:** Ekin Daplan, Luca Turin

## Abstract

Visible light triggers free radical production in alive and intact *Drosophila melanogaster*. We exposed fruit flies to red (613-631 nm), green (515-535 nm), and blue (455-475 nm) light while we monitored changes in unpaired electron content with an electron spin resonance spectrometer (ESR/EPR). The immediate response to light is a rapid increase in spin content lasting approximately 10 seconds followed by a slower, linear increase for approximately 170 seconds. When the light is turned off, the spin population promptly decays with a similar time course, though never fully returning to baseline. The magnitude and time course of the spin production depends on the wavelength of the light. Initially, we surmised that eumelanin might be responsible for the spin change because of its documented ability for visible light absorption and its highly stable free radical content. To explore this, we utilized different fruit fly strains with varying eumelanin content and clarified the relation of melanin types with the spin response. Our findings revealed that flies with darker cuticle have at least three-fold more unpaired electrons than flies with yellow cuticle. However, to our surprise, the increase in unpaired electron population by light was not drastically different amongst the genotypes. This suggests that light-induced free radical production may not exclusively rely on the presence of black melanin, but may instead be dependent on light effects on quinone-based cuticular polymers.

## Introduction

The absorption of visible light by matter is largely due to electronic transitions. Aside from a few highly coloured pigments and chromophores, molecular components of living things are mainly transparent to visible light. Visible light ranges in wavelength from 700 to 380 nanometers, from red to blue, corresponding to a photon energy of 1.8 to 3.3 electron volts (eV). The electronic band-gap of proteins and nucleic acids is of the order of 5eV (Albani 2011). Therefore, most of the contents of a fly are transparent to visible light. Furthermore, most of the molecules in living systems are closed-shell molecules hence invisible to ESR.

Drosophila melanogaster was chosen as a subject for our study because it is small enough to fit in an X-band ESR and is easy to produce and manipulate. Electron spin measurements in live flies were performed in our lab with ESR before (Turin, Skoulakis, and Horsfield 2014). ESR measurements with alive and intact animals larger than flies are not easy, partly because larger animals require L-band (≈1GHz) measurements to avoid intense microwave absorption by water, with attendant loss of sensitivity compared to X-band. Absorption of X-band microwaves by flies limits the number of flies to about 30. Several studies have been reported on the spin in living systems with ESR (Commoner and Ternberg 1961; Mallard and Kent 1966; Commoner, Townsend, and Pake 1954; Trapp et al. 1983).

The first barrier to visible light entering a fly is its cuticle. The composition and the chemical interactions that make the insect cuticle are not fully understood. However, it is known that the fly cuticle is layered (Nation and Sr. 2022). The outermost layers of the cuticle commonly contain lipids, specifically wax, in keeping with its purpose of water conservation. Deeper layers incorporate pigments, proteins, and carbohydrates, respectively. Chitin, one of the main biopolymers of the cuticle and the second most abundant polysaccharide on earth (Moussian 2013), is transparent to visible light (Neville 2012). The only known molecular category that exclusively provides color due to its chemical structure via visible light absorption in flies is melanin (Vásquez-Procopio, Rajpurohit, and Missirlis 2020). The other key polymer in flies is sclerotin. Sclerotins are known to function to provide structural strength (Pryor 1940). Interestingly, biochemical pathways of melanin and sclerotin intersect (Figure 2). While N-acetyl dopamine (NADA) sclerotin is reported to be colorless, N-β-alanine dopamine (NBAD) sclerotin exhibits yellowish hues (Wittkopp, True, and Carroll 2002).

Melanin is an amorphous polymer extensively produced by Drosophila melanogaster. The pigment is primarily located on the cuticle in the form of black (eumelanin) and brown melanin. The differences in the production and regulation of melanin types are partially known (Figure 2). Apart from the cuticular melanins, there is some evidence for the existence of molecular machinery to produce neuromelanin in drosophila (Barek, Veraksa, and Sugumaran 2018; Latocha et al. 2000). Neuromelanin is also a catecholamine-based pigment (Cao et al. 2021) that is found in some catecholaminergic nuclei of humans and some other mammals’ brains. Neuromelanin is well known for its association with Parkinson’s Disease. Accumulation of neuromelanin in the brain is considered to be age dependent (Zucca et al. 2023). Hence, for animals with a short life span like fruit flies, neuromelanin production seems to be absent.

Yellow and ebony genes play pivotal roles in determining the cuticle color of flies. The effects of the genes are easily observable in the phenotype (Figure 1). The genes are required in the synthesis of eumelanin (black melanin) and NBAD sclerotin (Figure 2). In the case of ebony knock-out flies, they are not able to form the link between dopamine and β-alanine (Hodgetts and Konopka 1973) to synthesize yellow sclerotin and have a very dark cuticle coloration. In yellow knock-out flies, the conversion of dopachrome into 5,6-dihydroxyindole for eumelanin production is hindered, leading to their distinctive yellow cuticle coloration.

**Figure 1.**
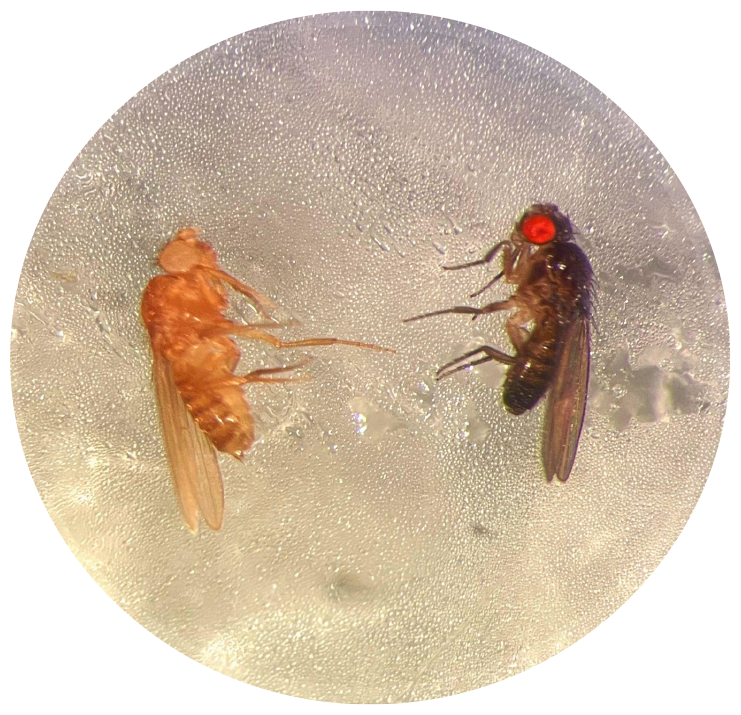
Different cuticle and eye colorization in Drosophila melanogaster. On the left is yw^1^ fly and on the right is e^1^.

**Figure 2.**
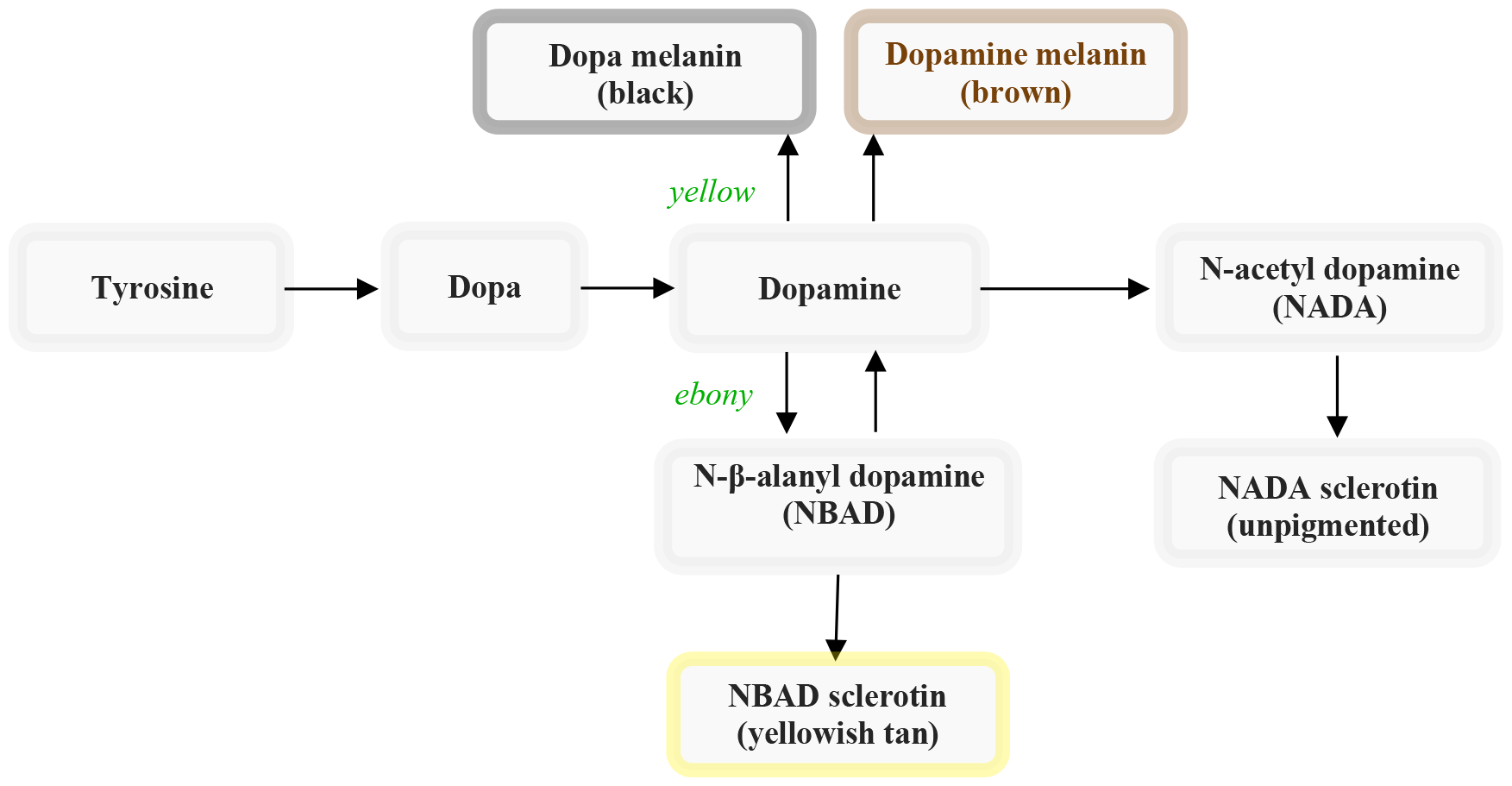
Melanin production pathways in flies. The figure is redrawn from (Massey and Wittkopp 2016).

The majority of ESR signals come from the cuticle. The ESR spectrum of fruit flies has been shown to depend on the presence of melanin and sclerotin (Kayser and Palivan 2006). The effect of light on both synthetic and natural melanin has been described previously in vitro (Arnaud et al. 1983; Sealy et al. 1984; Bibang et al. 1989; Mostert et al. 2018; Grieco, Kohl, and Kohler 2023). One of the studies (Mostert et al. 2018) hypothesized that the constituents of eumelanin comprised carbon-centered radicals and semiquinones, with semiquinones being implicated in the light response. Our objective was to investigate the light response of fruit flies with varying levels of black melanin content.

## Materials

### Drosophila melanogaster

w^1^(white), yw^1^(yellow, white), and e^1^(ebony), Canton-S, and Oregon-R strains, purchased from Bloomington Drosophila Stock Center were used. Strains are cultured on standard wheat-flour-sugar food supplemented with soy flour and CaCl2, at 25°C with 50–70% relative humidity in a 12 h light/dark cycle.

### LEDs

- Prolight-Opto LED Star Red 3W, 613.5-631 nm Typical Luminous Flux: 126 lm
- Prolight-Opto LED Star Green 3W, 515-535 nm Typical Luminous Flux: 197 lm
- Prolight-Opto LED Star Blue 3W, 455-475 nm Typical Luminous Flux: 55 lm

## Method

### Sample Preparation

ESR measurements of flies have previously been described by Turin et al (Turin and Skoulakis 2018). The ESR setup was modified to allow the LED light to reach the flies in the Teflon tube (Figure 3). In order to achieve this, a 3 cm long glass rod is inserted into the Teflon tube. 1 cm of the glass tube is nested inside the Teflon tube from one end. The glass tube, from the other end, is glued to the lens of the LED. The base of the LED is glued to a folded copper sheet for heat dissipation and easy handling. Finally, the LED is connected to a DC power supply.

**Figure 3.**
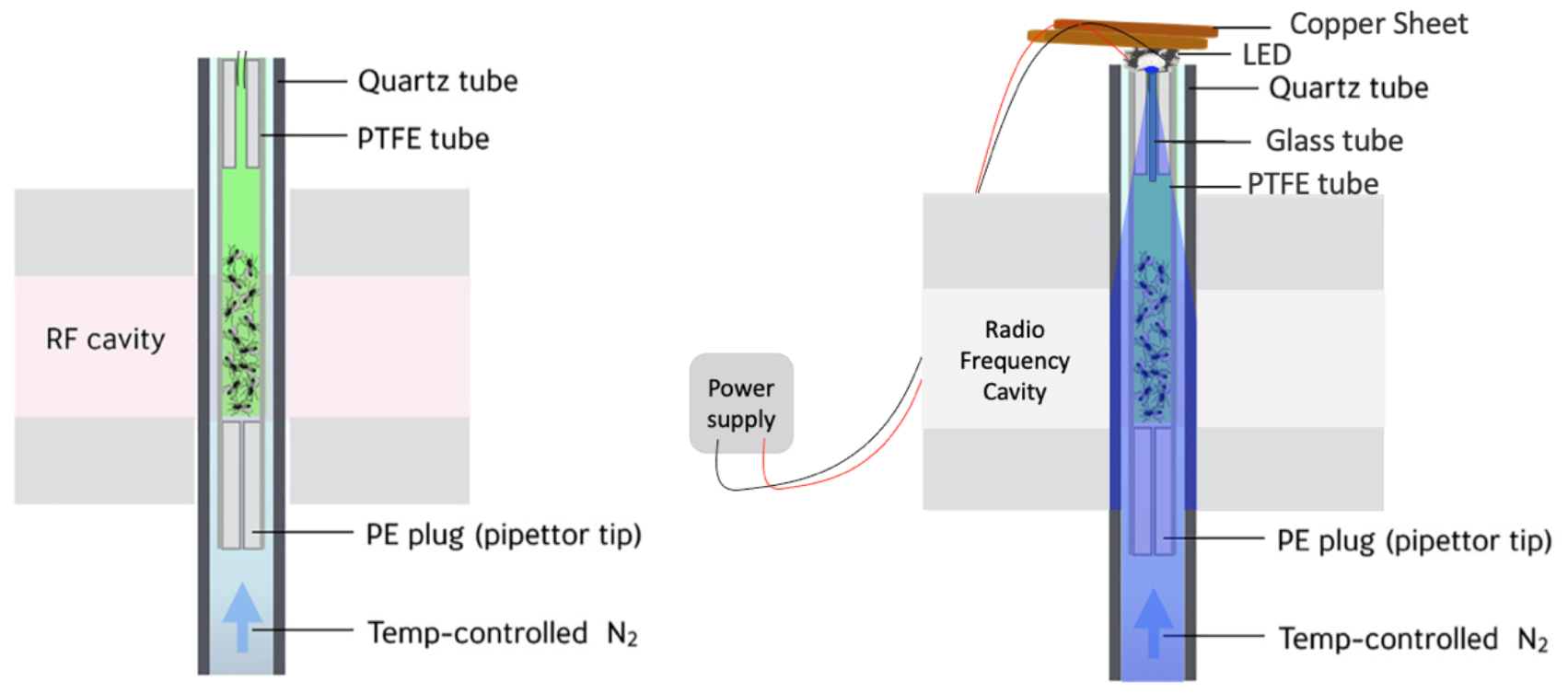
Left ESR spectrometer is a depiction taken from Turin et al.(Turin and Skoulakis 2018) Right Modified ESR spectrometer set up for the experiments of the paper.

### Experiment Design

Every experiment was done using 30 male fruit flies. The ESR was set to 9.4 GHz throughout the experiments. The ESR spectrum was collected with the same parameters for control and experimental groups. The central value was set to 3350 gauss with a 100 gauss sweep. All recordings were performed under ambient gas nitrogen to prevent fly movement without having to lower the ambient temperature. Drosophila melanogaster is known to survive anoxia up to 12 hours (Campbell, Werkhoven, and Harrison 2019).

Light response is measured under a stable magnetic field which is determined by the positive peak of the derivative curve of the ESR spectrum. Flies are recorded under a stable field for 120 seconds before 180 seconds of light exposure. After the light is turned off, the recording is sustained for an additional 300 seconds.

## Results

### 1. Light exposure produces a partly reversible increase in the population of unpaired electrons

In Figure 4, the graph on the left illustrates the standard ESR spectrum of thirty w^1^ flies. This spectrum is a single line, here observed in derivative mode, and it is centered around g=2.003. Upon exposure to blue light for three minutes there is an approximately ∼15% increase in spin, as depicted in the right panel of Figure 4. About half of this increment takes place during the first minute. After this initial rapid rise, the spin continues to increase in a more gradual, linear fashion. We can show this behavior quantitatively by fitting the data into a double exponential, represented by the black line in figure 4 on the left. At the end of the light exposure, turning off the LED causes ∼80% of the added spin to diminish within five minutes (Figure 4, right). The time constants observed during the light-off period resembles the light-on period. There is a substantial difference between the two time constants for the light-on and light-off periods, for both, τ_1_ being the very short one. Furthermore, both time constants shrink by about 1 second in light off period. The effect of blue light on the ESR spectrum is measurable even after 5 minutes through a magnetic sweep (Figure 4, left, blue trace), and it is calculated to be ∼5%.

**Figure 4.**
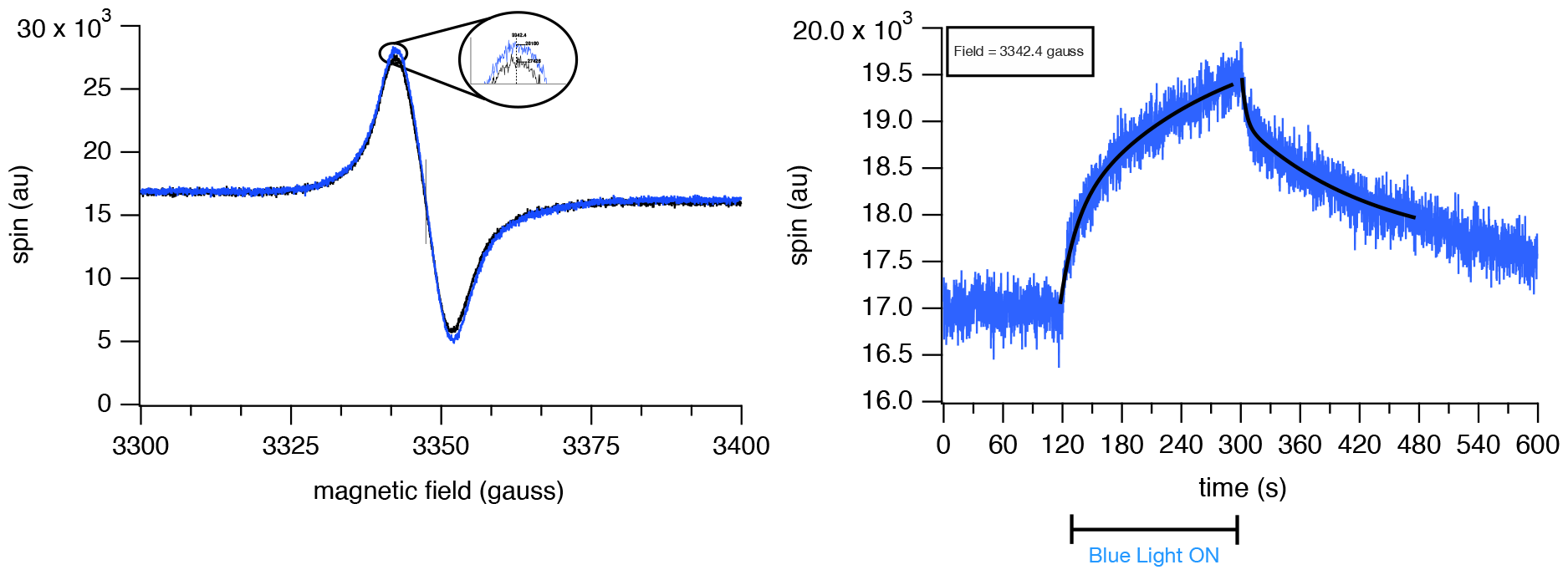
Exposure procedure applied to w^1^ flies with ambient gas nitrogen. Left, ESR spectrum of the sample before and after blue light. The black line is used to visualize the spectrum before light, and the blue color represents the spectrum after exposure. The relative spin before light exposure is 22090 arbitrary unit (au), calculated by subtracting the minimum from the maximum value in the black trace, and after exposure is 23570 au, therefore the increase is ∽1480 au with blue light exposure. Right, The spin response of the flies to blue light at a fixed value of magnetic field, 3342.4 gauss, corresponding to the positive peak of the derivative curve. The black lines show the double exponential fit. Time constants for light on trace are τ_1_=1.5 seconds, τ_2_=21.4 seconds and for light off trace τ_1_=0.7 seconds τ_2_=20.4 seconds.

### 2. Repeated light exposures cause further increases in spin

We applied the exposure procedure three times in sequence to the same group of w^1^ (wild type) flies (Figure 5). After these repeated exposures, we calculated the residual spin based on the ESR spectrum. The amount of retained spin was approximately twice that of a single exposure. The overall time course of the spin increase has been conserved with a slight increase in the peak value in each follow-up exposure (Figure 5, right). All the exposures fit well to the double exponential model.

**Figure 5.**
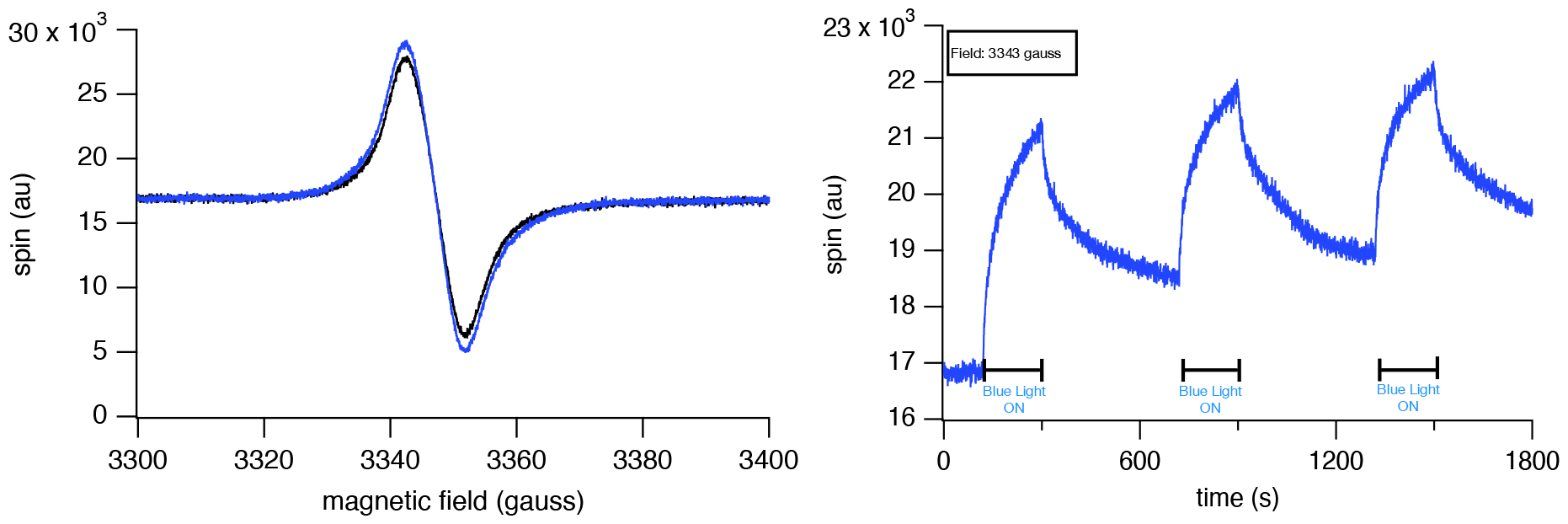
The spin response of the 2 different groups of flies to 3 consequent blue light exposure is shown in the figure. Left, The ESR spectrum of flies before (black) and after (blue) the light exposure is displayed. Right, Light response curve.

Although there was no gradual dramatic change amongst them, the difference between light-on and light-off periods for both τ_1_ and τ_2_ was minimal in the third exposure (Figure 6). Only for the light-on period, τ_2_ exhibited subtle, gradual shortening (Table 1). The difference between the light-on and light-off periods in both τ_1_ and τ_2_ was maximal in the second exposure, light-off period being the longer one.

**Figure 6.**
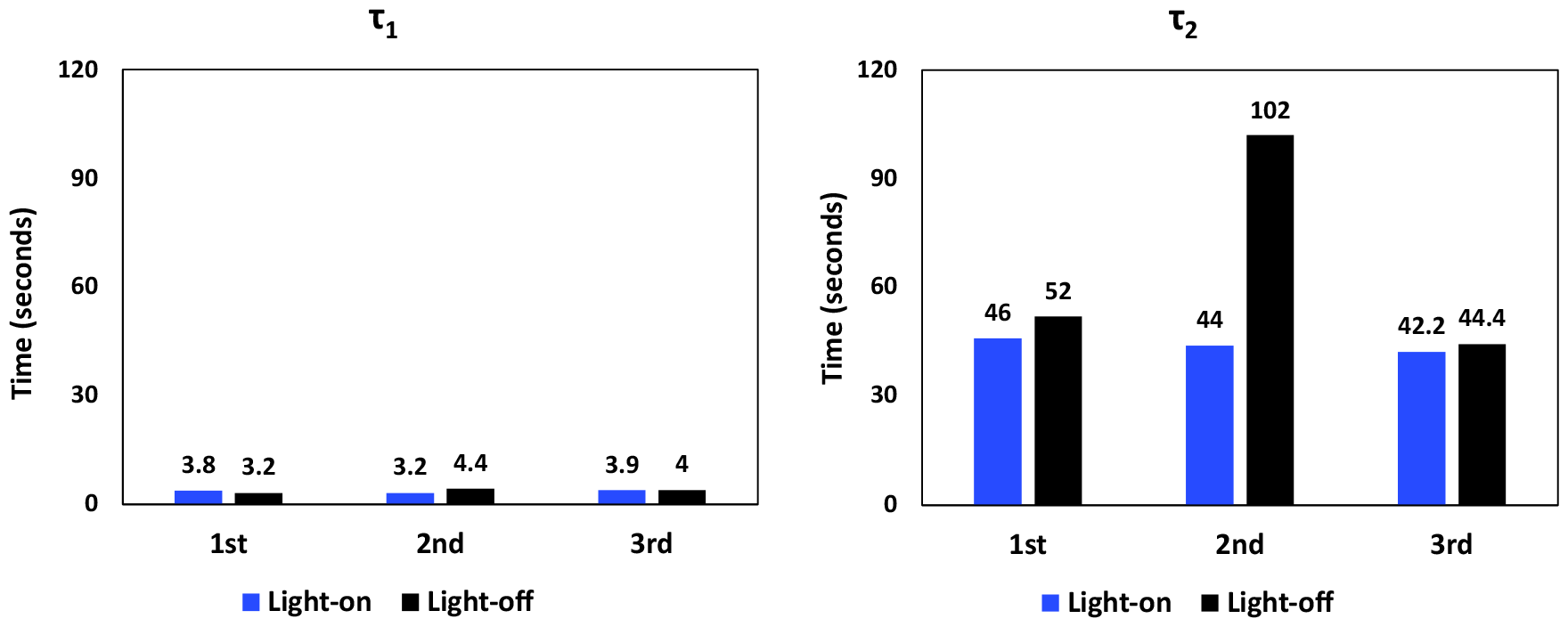
The graphs in seconds illustrate the change in τ_1_ and τ_2_ with successive exposures to blue light. Gray bars represent light-off and blue bars the light-on period.

**Table 1.** The table in seconds shows the change in τ_1_ and τ_2_ with successive exposures to blue light.

| time constant | exposure | light on | light off |
| --- | --- | --- | --- |
| $\tau_1$ | 1<sup>st</sup> | 3.8 | 3.2 |
|  | 2<sup>nd</sup> | 3.2 | 4.4 |
|  | 3<sup>rd</sup> | 3.9 | 4 |
| $\tau_2$ | 1<sup>st</sup> | 46 | 52 |
|  | 2<sup>nd</sup> | 44 | 102 |
|  | 3<sup>rd</sup> | 42.2 | 44.4 |

### 3. A short wavelength of light produces more spin than a long wavelength

After characterizing the signal with one wavelength, we wondered how different wavelengths alternate the light-induced spin response. For this purpose, we exposed different groups of w^1^ flies to red (613-631nm) and green (515-535nm) in addition to blue (455-475nm) light. Short-wavelength causes higher spin production in comparison to long-wavelength (Figure 7, right). The exponential increase that takes place within the first 25 seconds displayed a similar magnitude for red and green LEDs, whereas the blue LED caused a larger increase. These differences cannot be ascribed to light output per se, since the light flux from the blue LED was approximately three times smaller than from the green and comparable to the red LED output.

**Figure 7.**
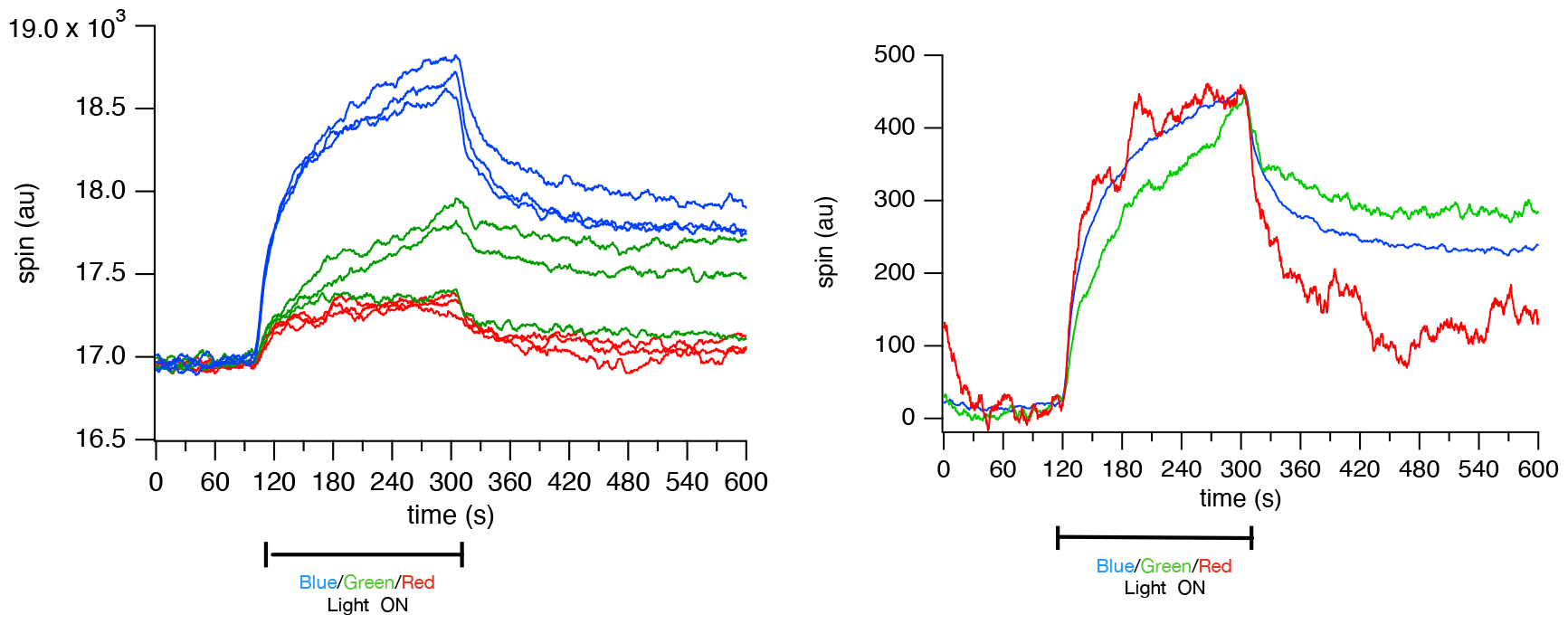
Left The spin responses of the flies to the different colors of visible light are displayed. The colors of the lines indicate the response to the corresponding wavelength of light exposure. The peak value of the ESR spectrum trace is picked for the stable magnetic field measurements. Accordingly, the magnetic field was ∼3342 gauss for blue, ∼3343 for green and ∼3343 for red light assessment. Right Shows the averaged and scaled form of the graph on the left.

To study differences in the kinetics of the response to various light wavelengths, we scaled the traces in respect to their peak values. We observed a difference in the response to green light when compared to red and blue light. Red and blue traces fit well a double exponential model, with similar time constants. However, the first time constant of green light was slightly longer, and the second relatively larger (τ_1_ /τ_2_ was around 1.5/12 for red and blue and 2.9/120 for green light). None of our measurements displayed as long τ_2_ as the green light. The light-off period revealed other differences. Red light exhibited a longer τ_2_, corresponding to 48 seconds, while green and blue trace had a much shorter τ_2_, blue τ_2_ =9 seconds, green τ_2_ =12. The deviations in the time constants between different light exposures seem to be less in τ_1_ and mostly in τ_2_ (Table 2).

**Table 2.** The table in seconds illustrates the change in τ_1_ and τ_2_ with exposures to blue, green and red light.

| time constant | Color | light on | light off |
| --- | --- | --- | --- |
| $\tau_1$ | Red | 1.1 | 1.6 |
|  | Green | 2.9 | 1 |
|  | Blue | 1.8 | 1.5 |
| $\tau_2$ | Red | 8.8 | 48 |
|  | Green | 120 | 12 |
|  | Blue | 14 | 8.8 |

### 4. Flies with different melanin contents have different baseline spin

The connection between black melanin production in fruit flies and its impact on the ESR spectrum has been reported before (Turin and Skoulakis 2018). Here, we confirm this, demonstrating that the absence of black melanin leads to a reduction in the ESR spectrum signal (Figure 8). On the other hand, the absence of yellow sclerotin is associated with darker flies and an increased intensity in the spectrum (Figure 8, black line). We can quantitatively interpret the intensities by considering the relationship of peak values, where yw^1^ flies exhibit approximately ⅓ of w^1^ and ¼ of the spin intensity of e^1^ flies.

**Figure 8.**
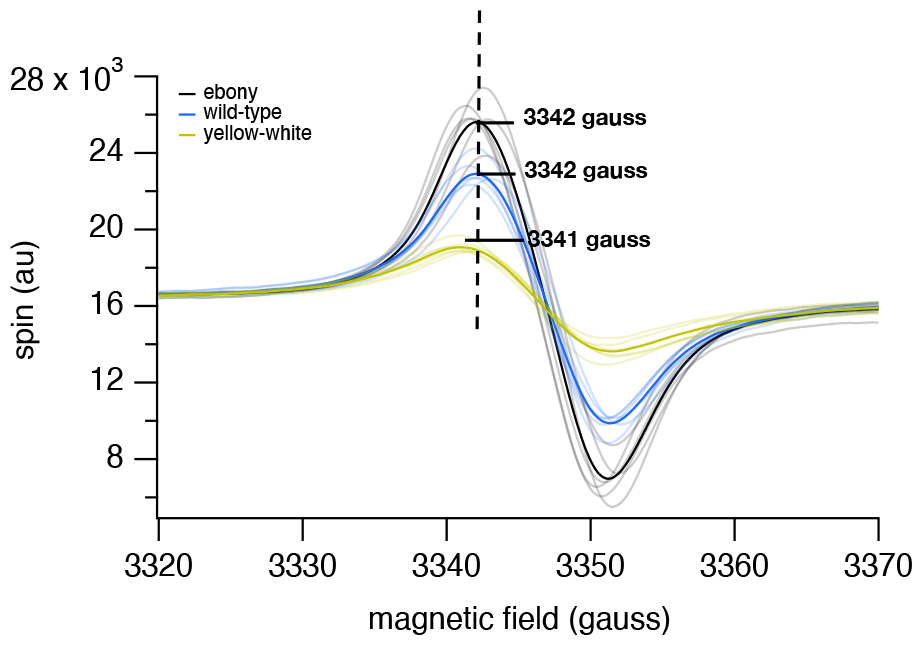
The plot shows the ESR spectrum of flies with different eumelanin and yellow sclerotin content. Black lines are the measurements of e^1^ (ebony) flies, blue are w^1^ (wild type), and yellow are yw^1^. Average ESR spectrum of each genotype was calculated from the measurements of six different groups of flies per each genotype, represented by the respective colors in the graph in bold. The maximum values observed in the traces are 25614 for black, 22902 for blue and 19075 for yellow. Both black and blue traces are aligned at the same magnetic field for their peak values, 3342 gauss. However, the yellow trace is slightly shifted left and has a peak at 3341 gauss.

### 5. Relative light-induced spin production is higher in yw^1^ flies

After measuring the ESR spectra for e^1^, w^1^, and yw^1^ fruit flies we examined their light responses. Since eumelanin is repeatedly associated with photoprotective features and presented with high intensity ESR spectrum, we expected a relatively strong response. We anticipated correlation with the baseline spin, i.e. a lower light-induced spin production by yw^1^ flies and higher light-induced spin production by ebony. Strangely, however, yw^1^ and e^1^ flies showed only a slight difference, with more spin increase in yw^1^ flies under illumination (Figure 9, left).

**Figure 9.**
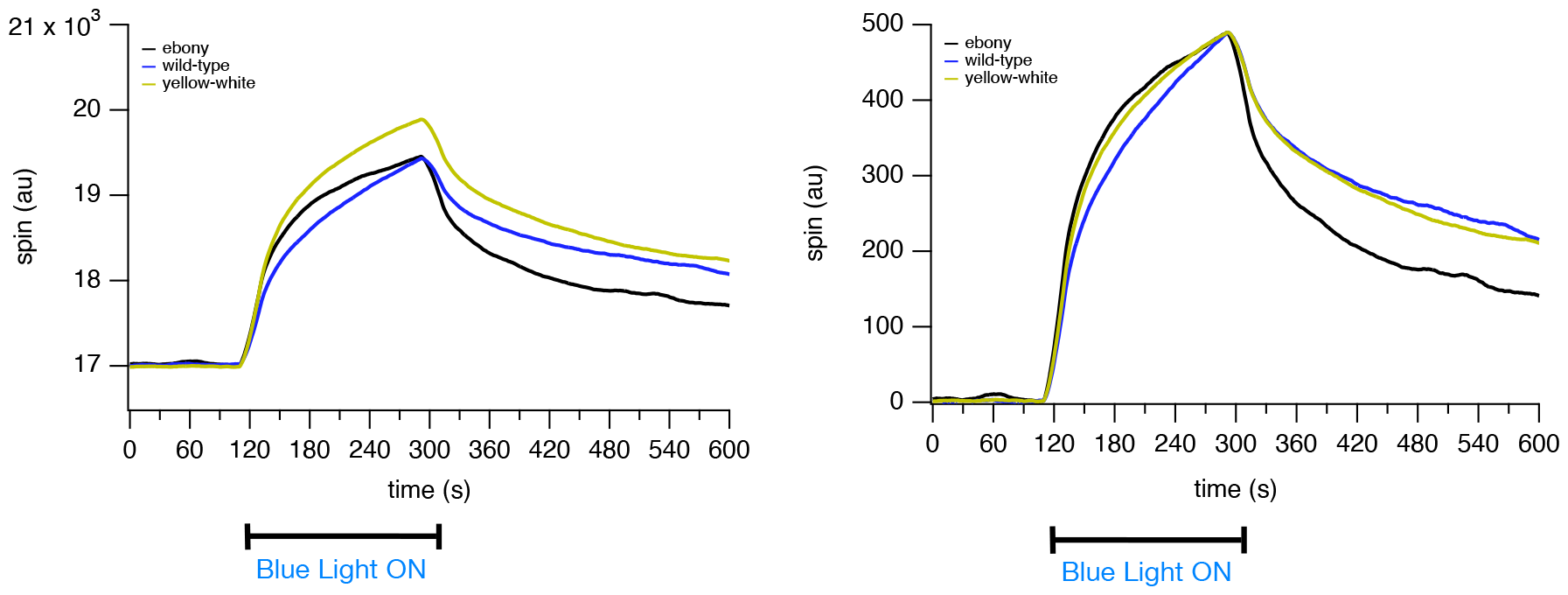
Left The light response of w^1^ (blue) e^1^ (black) and yw ^1^ (yellow) flies are displayed. The average of 6 distinct fly groups of each genotype is plotted with a marker. Right The average of the light response curves are scaled at the peak value. The magnetic field value was stable at 3342 gauss for e^1^ and w^1^, and 3341 gauss for yw^1^ flies.

The constant τ_1_ showed a gradual decrease towards the darker flies. For both light on and light off period τ_1_ was shortest in ebony flies in comparison to others (Figure 10, left). Also, the difference between the two light periods was maximum in ebony flies. On the other hand, τ_2_ did not display a similar pattern between the genotypes. The only disparity was that ebony flies had very similar τ_2_ for both light on and off periods, unlike yellow-white or white (Table 3).

**Figure 10.**
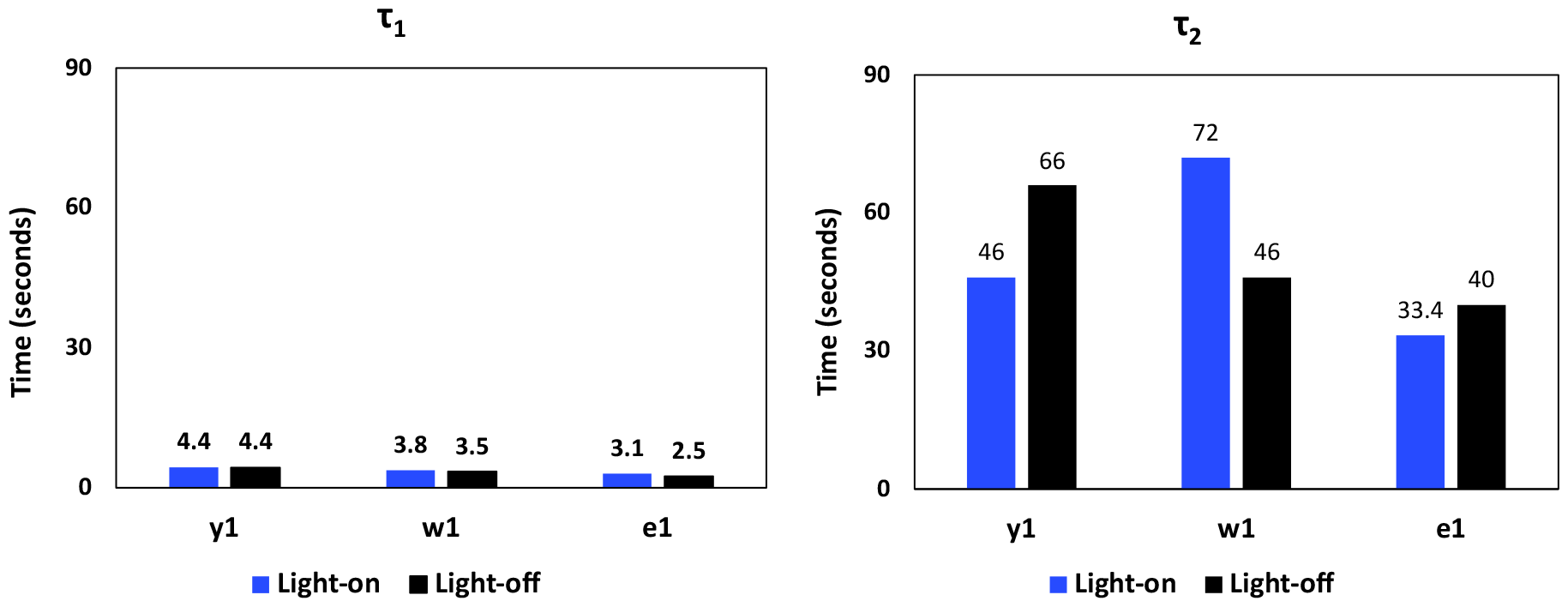
The graphs in seconds illustrate the change in τ_1_ and τ_2_ with exposure of the three mutants to blue light. Gray bars represent light-off and blue bars the light-on period.

**Table 3.** The table in seconds illustrates the change in τ_1_ and τ_2_ with exposures to blue light in yellow-white, white, and ebony fruit flies.

| Time Constant | Genotype | Light-on | Light-off |
| --- | --- | --- | --- |
| $\tau_1$ | $y^1$ | 4.4 | 4.4 |
| | $w^1$ | 3.8 | 3.5 |
| | $e^1$ | 3.1 | 2.5 |
| $\tau_2$ | $y^1$ | 46 | 66 |
| | $w^1$ | 72 | 46 |
| | $e^1$ | 33.4 | 40 |

## Discussion

Exposing intact and alive Drosophila melanogaster to visible light results in an increase in the population of free radicals. This increase can be monitored with ESR directly. Initially, we predicted pigments to be the cause of the increase in free radicals due to their light absorption properties. The ESR spectrum of differently pigmented fruit flies has been described before (Turin and Skoulakis 2018). Although complete constituents yielding the ESR spectrum of fruit flies are not yet elucidated, the ESR signal clearly follows melanin content. We therefore focused especially on melanin. Several studies have reported the free radical production caused by visible light by synthetic melanin (Sealy et al. 1984; Sarna and Sealy 1984; Sever, Cope, and Polis 1962; Pascutti and Ito 1992; Kalyanaraman, Felix, and Sealy 1984; Mostert et al. 2018). In only one study no change in ESR signal during red light exposure to melanin was reported (Mostert et al. 2012). As a first approximation, we measured the light response of flies that use different melanization pathways, w^1^, yw^1^, and ebony (Wittkopp, Carroll, and Kopp 2003). We expected ebony mutants to exhibit a larger response to light. To our surprise, we couldn’t find any significant difference between the dark ebony and the light-colored yellow mutants (Figure 9). In fact, yellow mutants exhibit a slightly larger increase in response to light.

This raises the question of what constituent of cuticle is visible in ESR and has the required optical characteristics to contribute to the spin response to light on flies. The stable free radical content of the cuticle, in the absence of melanin, is reported only in one study with ESR (Kayser and Palivan 2006). It has been proposed that semiquinones to be the moiety within eumelanin that causes an increase in the ESR signal due to light exposure (Mostert et al. 2018). Therefore, two additional candidates could account for light-induced spin increases beside brown melanin (dopamine melanin): semiquinone-like melanin precursors and yellow sclerotin. Sclerotin cannot be the only element to give rise to the signal, since it is absent in ebony flies. There is not much information concerning the presence and regulation of brown melanin. Considering the intricate relationship of these chemical constructs in their biosynthesis (Figure 2), a more reasonable assertion would be that various types of melanin, their constituents, semiquinones, and similar molecules like sclerotin collectively add up to the response to visible light.

The overall light response characteristics such as following a double exponential with no lag are conserved across different wavelengths of light and various fly groups (Figure 7, Figure 9) with small variations in the magnitude. Repeated exposures showed that at every light exposure the flies responded in a similar manner (Figure 5) with a slight reduction in the amplitude of the response increase. This suggests a limitation in the sample’s ability to generate free radicals upon continued light exposure. We observed another interesting difference in the traces while experimenting with three colors of LED. It is plausible that the generation of light-induced free radicals in flies follows a linear trend, considering eumelanin’s absorption spectrum increases through decrease in the wavelength of light (Meredith et al. 2006). However, our result (Figure 7) revealed that the response to green light did not conform to this expectation.

The structure and function of melanin types still lack a complete understanding (Cao et al. 2021). In this paper we propose a method to investigate the properties of melanin and to produce electron spin in vivo which may be of use to further genetic and photochemical studies.

## Acknowledgments

The authors gratefully acknowledge support from Ionis Pharmaceuticals and assistance from ADVIN Solutions.

